# gpps: An ILP-based approach for inferring cancer progression with mutation losses from single cell data

**DOI:** 10.1101/365635

**Authors:** Simone Ciccolella, Mauricio Soto Gomez, Murray Patterson, Gianluca Della Vedova, Iman Hajirasouliha, Paola Bonizzoni

## Abstract

**Motivation:** In recent years, the well-known Infinite Sites Assumption (ISA) has been a fundamental feature of computational methods devised for reconstructing tumor phylogenies and inferring cancer progression where mutations are accumulated through histories. However, some recent studies leveraging Single Cell Sequencing (SCS) techniques have shown evidence of mutation losses in several tumor samples [19], making the inference problem harder.

**Results:** We present a new tool, `gpps`, that reconstructs a tumor phylogeny from single cell data, allowing each mutation to be lost at most a fixed number of times.

**Availability:** The General Parsimony Phylogeny from Single cell (`gpps`) tool is open source and available at https://github.com/AlgoLab/gppf.

## 1 Introduction

Recent developments of targeted therapies for cancer patients rely on the accurate inference of the clonal evolution and progression of the particular cancer. As discussed in several recent studies, such as [24] and [32], understanding the order of accumulation and prevalence of somatic mutations during cancer progression can help better devise therapeutic strategies. Moreover, studying the evolutionary history of tumors can provide some insights on which mutations lead to drug resistance.

The most widely studied techniques for inferring cancer progression rely on data from next-generation bulk sequencing experiments. In these cases, we sample mixtures of cells that are not homogeneous from a mutational profile (*i.e.*, which mutations appear in a cell) point of view. Moreover, we cannot easily distinguish between cells: the only information we can have is, for each mutation, the fraction of cells in a sample carrying such mutation. Recently, many computational approaches have been developed for the analysis of bulk-sequencing data with the purpose of inferring tumoral subclonal decomposition and reconstructing tumor phylogenies (trees) [3, 9, 15, 17, 22, 23, 25, 30, 31, 33], but almost all of them model a tumor progression as the accumulation of mutations under the Infinite Sites Assumption, that is recurrent mutations and mutation losses are not allowed.

Single Cell Sequencing (SCS) greatly improves the resolution of the data available, as it provides the set of mutations of each cell analyzed. However, this technique is currently expensive and not especially reliable, since it produces datasets with a high amount of noise that include allelic dropout (false negatives) and missing values, due to lack of read coverage, as well as false positive calls – although this event is much rarer. Another source of noise is due to doublets, that is signals originating from two separate cells which are erroneously inferred to originate from a single cell: we point out this latter problem is fading away and can be tackled computationally. Still, we need efficient methods that are able to cope with the kind of data that SCS techniques are currently producing, by taming the difficulties due to the noise in data.

Various methods have been developed for this purpose [16, 28, 34], some of them introducing a hybrid approach of combining both SCS and VAF data [20, 21, 26, 29]. As stated before, most of these methods rely on the Infinite Sites Assumption (ISA), which states that a mutation is acquired at most once in the phylogeny and is never lost. The ISA was introduced in [18]. This simplifying assumption also leads to a computationally tractable model of evolution [11] called the perfect phylogeny. Cancer progression, however, is a very fast and aggressive form of evolution with limited data supporting neutral evolution [6], with some studies showing rather the evidence of selection [2, 6] – something that is particularly true in tumor samples after a relapse [6, 8, 10], where the tumor has already been highly selected by the therapy targeted to destroy it. Thus, one would be expect that we must abandon the strict Infinite Sites Assumption in this setting, and indeed this is the case, as some recent studies are finding hints suggesting that the ISA does not always hold [2, 4, 19]. In [4], the authors find that large deletions on several branches of a tree can span a shared locus, thus a given mutation may be deleted independently multiple times. In [2], the authors show that in certain cases, homozygous deletions in cancer genomes can even provide a selective growth advantage. Each (independent) deletion of an acquired mutation takes us further away from the ISA. Some recent methods such as TRaIT [26] and SiFit [34] allow deletions of mutations.

The Dollo model [27] of evolution is designed exactly for some of the cases where a perfect phylogeny does not represent the actual data. More precisely, the Dollo model requires each mutation to be acquired exactly once in the entire history analyzed, while removing all restrictions on the number of times that a mutation can be lost. The Dollo model as well as the Dollo(*k*) variants, where each mutation can be lost at most *k* times, has been introduced recently in the literature on algorithmic approaches for tumor progression inference [3, 5]. Unfortunately, the Dollo model does not have the convenient computational tractability of the perfect phylogeny model [11], hence requiring more sophisticated algorithms.

In this paper we propose (`gpps`), a tool of the gppf family, that exploits an Integer Linear Programming (ILP) approach to infer a tumor progression that can include mutation losses, from single cell sequencing data.

**Figure 1:**
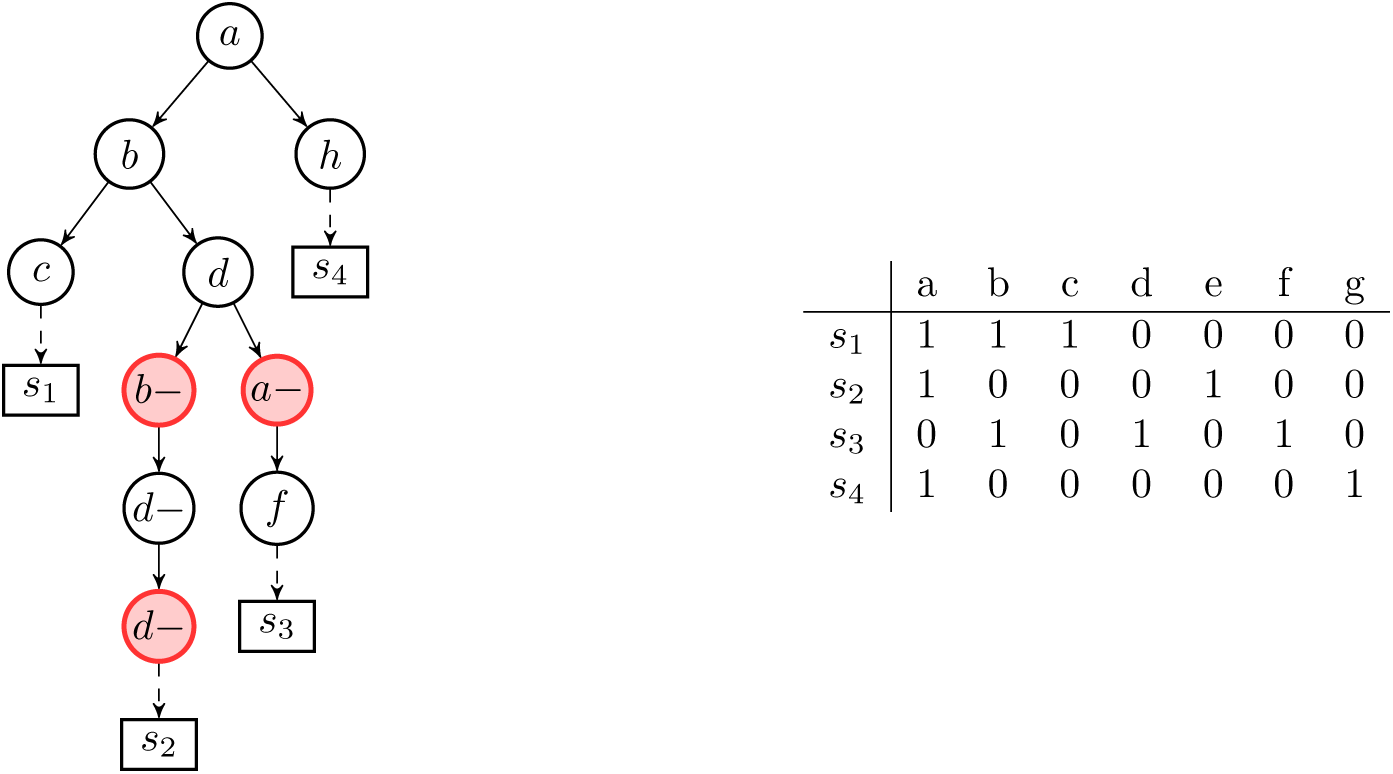
Example of a binary matrix that does not allow a perfect phylogeny, since columns *a* and *b* are in conflict, *i.e.*, the four gametes rule [11] does not hold. The tree represents one of the possible Dollo phylogenies that explain the matrix.

## 2 Tumor phylogeny reconstruction from single cell data

In the most abstract formulation, we can see the cancer progression reconstruction problem as a character-based phylogeny reconstruction problem [12] where each character represents the presence/absence of a specific mutation in a cell.

The input of the problem is an incomplete binary matrix *I*, where the entry *I*[*c, m*] = 0 indicates that the cell *c* does not have the mutation *m*, while *I*[*c, m*] = 1 indicates that the cell *c* has the mutation *m*. Finally, we denote with *I*[*c, m*] = ? where there is not enough information on the presence/absence of mutation *m* in cell *c*. We recall that uncertainty about the presence of a mutation in a cell is a consequence of insufficient coverage in the sequencing, hence it is unavoidable.

However, uncertainty is not the only issue in the sequencing process: the input matrix *I* also contains false positives and false negatives. We assume that these errors occur independently and uniformly across all the (known) entries of *I*. Namely, *P* denotes the predicted matrix, *i.e.*, the binary matrix without missing values computed by the algorithm. In this case, *α* denotes the false negative rate and *β* denotes the false positive rate. In other words, for each pair (*c, m*),

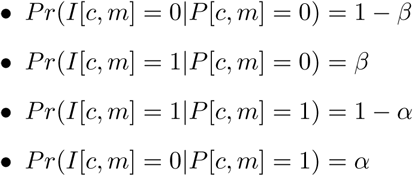

**Figure 2:**
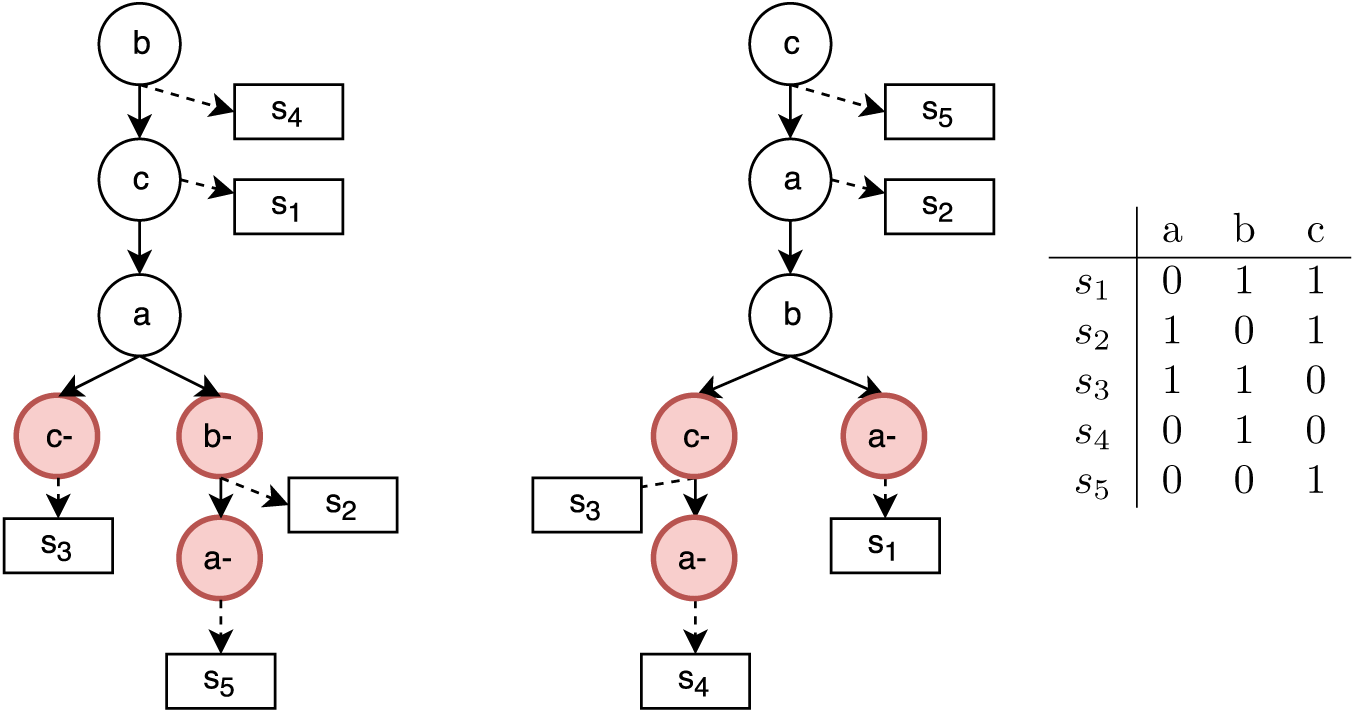
Example of two Dollo phylogenies that explain the same binary matrix. It is important to notice that the ancestral order of mutations *c, a* and *b* is inverted but the two different trees can equally explain the input binary matrix. In fact, in a Dollo phylogeny the order of two mutations can be inverted and, thank to the introduction of deletions, they could both be correct solutions for a given input.

Our goal is to find a matrix *P* that (1) corresponds to a phylogeny on the set of cells, and (2) maximizes the the likelihood

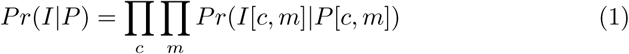

of the observed matrix *I* [16]. In other words, we want to find the phylogeny, as expressed by the matrix *P*, that maximizes the likelihood of the observed matrix *I* [16]. We point out that the values of the unknown entries of the input matrix do not factor into the objective function.

A phylogeny is a rooted labeled tree *T*, where the label set corresponds to the set of mutation gains and losses. The state *S*(*x*) of a leaf *x* in *T* is defined as the set of mutations that are acquired and not lost in the path from the root of *T* to *x*. We say that the tree *T* encodes a matrix *P* if there exists a mapping *σ* of the rows of *P* to the leaves of *T* such that for each row *r* of *P*, it follows that *C*(*r*) = *S*(*σ*(*r*)) where *C*(*r*) is the set of columns which are 1 in *r*, and *σ*(*r*) denotes the leaf of *T* associated with *r* through *σ*. In other words, in the tree *T* we assume that the cell *c* has been extracted from the subpopulation *σ*(*c*). See Figure 1 for example of a phylogeny a matrix that it encodes.

We can express the likelihood of the matrix *P* as in Equation 1 — since the involved probabilities are in [0,1] it is convenient to move to a (linear) log-likelihood maximization objective function of the form:

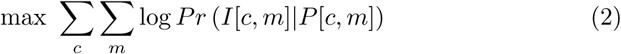

### 2.1 The evolution model

The Dollo parsimony rule can be interpreted as the impossibility of having an identical mutation in the evolutionary trajectory. This rule can be translated in the phylogeny tree model as the unique introduction of any single mutation but any number of deletions of this mutation.

From an algorithmic point of view, phylogeny reconstruction with a Dollo evolutionary model is an NP-complete problem [1, 7]. A hierarchical chain of restricted versions of the model can be obtained by bounding the number of deletions for each character. We denote as Dollo(*k*) the evolutionary model in which each mutation can be acquired exactly once and can be lost at most *k* times. In this way Dollo(0) and Dollo(1) correspond to the perfect and persistent phylogeny models, respectively. In the tree generation process for the Dollo(*k*) model (*k* > 0) we are required to augment a perfect phylogeny representing the cancer progression by adding nodes which represent the loss of a mutation, *i.e.*, a node labeled 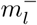, representing the *l*-th loss of mutation *m*. The state of the leaf *x* is the set of mutations *m* that, in the path from the root to *x*, have been acquired — the path has a vertex labeled *m*^+^ — but never lost — the path has no vertex labeled 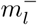. We stress that, when deletions are introduced, the set of feasible phylogenies which represent a given solution is no longer unique as in the case of perfect phylogeny — See Figure 2 for an example.

Testing if an incomplete matrix *I* has a phylogeny under the Dollo(*k*) model has been attacked via ILP for *k* = 0 [14], *k* = 1 [13], and for general *k* [3]. We will exploit the latter formulation to describe an ILP approach for tumor phylogeny reconstruction from single cell data.

First, we recall that a well known characterization of perfect phylogenies states that a *complete* binary matrix *M* has a directed perfect phylogeny if and only if it has no *conflicting* pair of columns — two columns are in conflict if they contain all three configurations (0, 1), (1, 0), (1, 1) — inducing the so-called forbidden matrix (*c.f.* Figure 1).

The ILP formulation on *incomplete* matrices [14] essentially consists of introducing a binary variable for each missing entry, and describing a set of constraints towards the goal of minimizing the conflicting pairs.

To adapt this approach to persistent phylogenies [13], we need a property:

**Proposition 1.** *[3] Let M be an incomplete binary matrix. Let M*_*e*_ *be the (incomplete) matrix obtained from M as follows: for each entry M* [*i, j*] *we have k* + 1 *entries M*_*e*_[*i, j*^+^] *and* 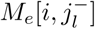 *(for* 1 ≤ *l* ≤ *k) such that (1) if M* [*i, j*] = 1 *then M*_*e*_[*i, j*^+^] = 1 *and* 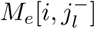 = 0 *for* 1 ≤ *l ≤ k, (2) if M* [*i, j*] = 0 *then the entries M*_*e*_[*i, j*^+^], 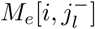*are all missing. Then M has a completion M*^*\**^ *that has a Dollo*(*k*) *phylogeny if and only if M*_*e*_ *has a completion* 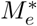 *such that if M* (*i, j*) = 0 *then* 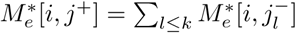*See Figure 3 for an illustration.*

Our main contribution is combining the ILP formulation of [3] with the definition of tumor perfect phylogeny reconstruction from single cell data, to obtain an ILP approach for tumor phylogeny reconstruction from single cell data that incorporates mutation losses in the model.

**Figure 3:**
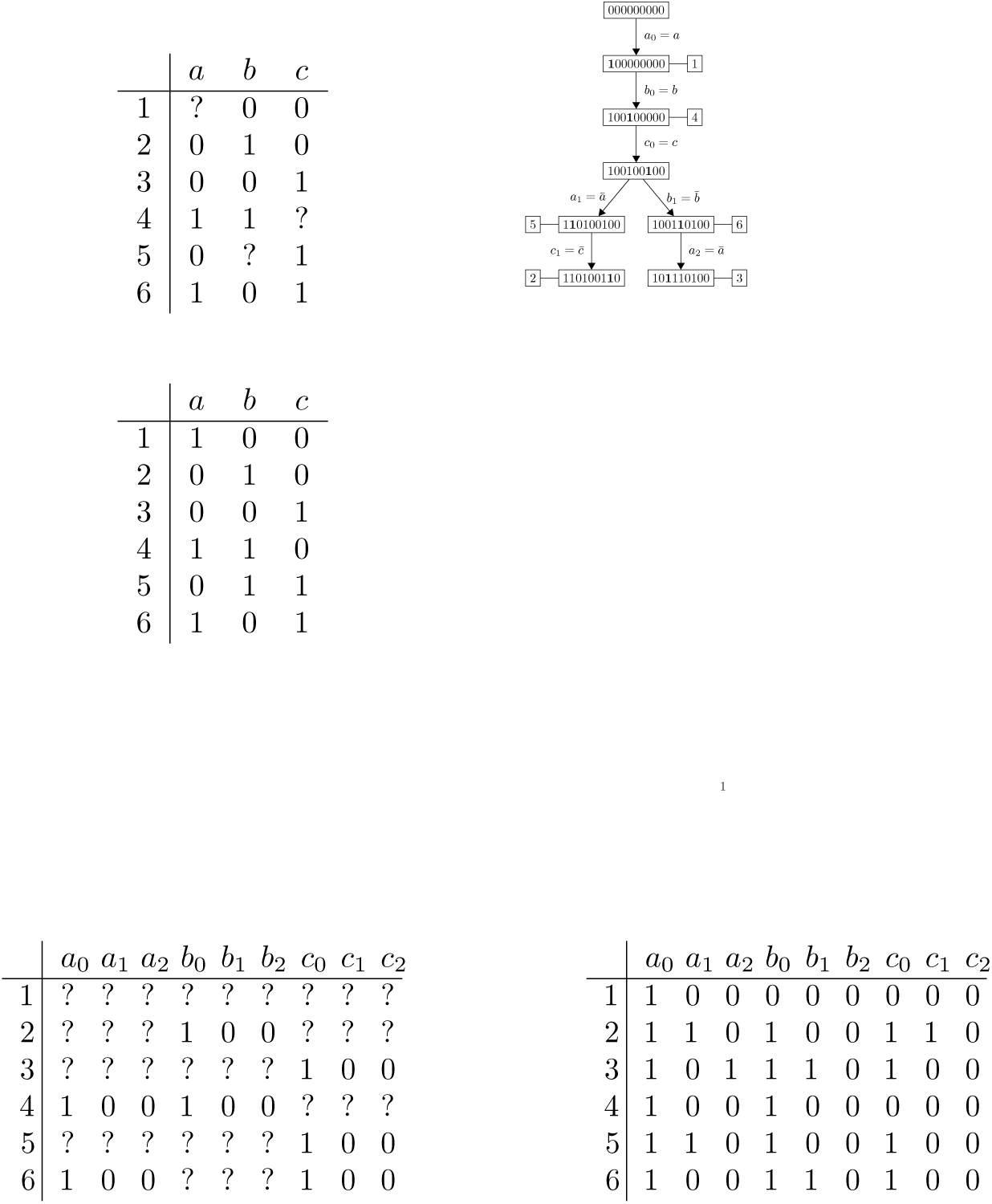
An input matrix *M* (top left), a Dollo(2) completion *M*_*c*_ (center left) and its corresponding phylogeny tree *T* (top right). The *M*_*D*(2)_ extended matrix (bottom left) and a completion for the MIDPP(*M*_*D*(2)_, *R*_*D*(2)_) according to Proposition 1. In the tree, boldfaced character corresponds to changes between each node and its parent.

## 3 The ILP formulation

In this section we present our ILP formulation for the tumor phylogeny reconstruction from single cell data.

We recall that the input of the problem is an incomplete matrix *I* represented as a set of binary variables *I*(*c, m*) such that *I*(*c, m*) = 1 if cell *c* has (according to the input data) the mutation *m*, while *I*(*c, m*) = 0 if cell *c* does not have (according to the input data) the mutation *m*. Notice that the input data is incomplete, hence we can have pairs (*c, m*) such that the variable *I*(*c, m*) does not exist.

The variables *P* (*c, m*^+^) and 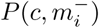 encode the extended matrix that we want to compute and that will satisfy Proposition 1. Differently from the variable *I*(.,.), for each pair (*c, m*), all variables *P* (*c, m*^+^) and 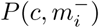exist.

We introduce some auxiliary variables that help in making the ILP formulation easier to read. The binary variables *F* (*c, m*) indicates if, in the predicted matrix, the cell *c* has the mutation *m*. By Proposition 1, *F* (*c, m*) = 1 if and only if *P* (*c, m*^+^) = 1 and all 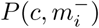are equal to zero. Moreover, the real variables *w*(*c, m*) represent the probability of *P* (*c, m*) given *I*(*c, m*) — the formula of the actual values depends on the possible cases, that is if we have a true positive, a true negative, a false positive, and a false negative.

To establish if two columns are in conflict, we introduce the final binary variables *B*(*p, q, a, b*), which are defined for each pair of columns (*p, q*) and for each possible pair of values (*a, b*)∈{(0, 1), (1, 0), (1, 1)}. More precisely, *B*(*p, q, a, b*) indicates if for the pair (*p, q*) of columns there exists a cell *c* where *P* (*c, p*) = *a* and *P* (*c, q*) = *b*. Notice that two columns *p* and *q* are conflicting iff *B*(*p, q,* 0, 1) + *B*(*p, q,* 1, 0) + *B*(*p, q,* 1, 1) = 3. We are now ready to introduce our ILP formulation, where we use *C* to denote the set of cells (*i.e.*, the rows of the input matrix *I*), *M* to denote the mutations (*i.e.*, the columns of *I*), and *M*^*\**^ to denote the set of possible mutation gains or losses.

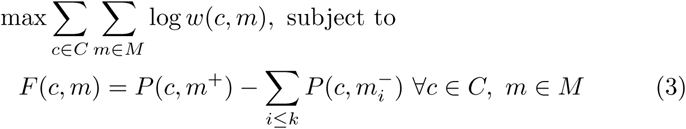

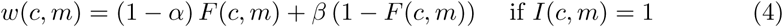

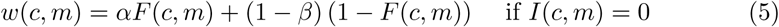

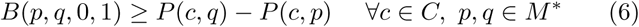

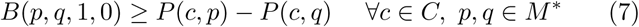

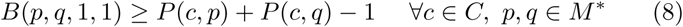

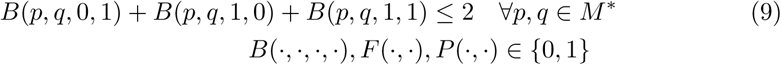

The total number of variables and constraints in the formulation are *O*(*nm* + *m*^2^) and *O*(*nm*^2^) respectively.

### 3.1 ILP implementation: gpps

Our approach has been implemented with a Python program called `gpps`. The code, data and scripts used in our experimental analysis is available at https://github.com/AlgoLab/gppf. The algorithm receives as input a frequency matrix *F*, the evolution model (persistent, Dollo(*k*), Camin-Sokal(*k*)) to be considered and the maximum number of clones in the clonal matrix (expressed as the percentage of the total number of mutations). The program generates the ILP formulation which is fed to an ILP solver in order to get the optimal solution.

In our experiments we have used Gurobi 6.5.2 as the ILP solver. Moreover, from the computed solution the program can construct the corresponding solution tree, provided that feasible solution has been found. Additionally, we have introduced a timeout on the running time, since the generated ILP problem could be large and its resolution could require a considerable amount of time. We exploit the fact that Gurobi can be halted at any time and it returns the best feasible solution computed so far. Hence, imposing a timeout allows the ILP solver to compute a solution with a small total error.

## Acknowledgement

SC acknowledges a Mobility Exchange Fellowship from the University of Milano-Bicocca. Part of this work has been done during SC visit at Weill Cornell. This work was also supported by start up funds (Weill Cornell Medicine) to IH.

